# Mapping the brain basis of appraisals and discrete emotions

**DOI:** 10.64898/2026.08.17.745188

**Authors:** Qingying Ye, Severi Santavirta, Asli Erdemli, Jinglu Chen, Vesa Putkinen, David Sander, Lauri Nummenmaa

## Abstract

The Component Process Model of Emotion conceptualizes any emotional episode (e.g., the discrete emotions of sadness, anger, fear, or interest) as being driven by the multiple appraisal components. However, both the specificity of the neural mechanisms underlying appraisal processes and the way these appraisal networks relate to the neural circuits underlying discrete emotions remain unclear. Here we investigated the neural correlates of appraisal processes and compared them with those of discrete emotions. Participants (n = 97) were scanned with functional magnetic resonance imaging (fMRI) while watching short movie clips with varying emotional contents. Intensity for 12 appraisals and 12 basic and epistemic emotions evoked by the movie clips were rated by independent participants (n = 444). The neural responses were modelled with convolved ratings of appraisals and discrete emotions. The results indicated that appraisals and discrete emotions are supported by a shared set of distributed brain regions that extend beyond typically reported emotion-related areas, encompassing perceptual, action-related, and higher-order cognitive systems. Activations were more consistent for and better explained by appraisals versus discrete emotions. Within this network, epistemic emotions elicited less consistent activations than basic emotions, particularly in limbic regions. Our results highlight the functional organization of appraisals and discrete emotions under dynamic and complex conditions and indicate that appraisal theories better explain neural responses than discrete emotion models.

## 1 Introduction

Emotions are evolutionarily conserved mechanisms that guide adaptive behavior, threat avoidance, and opportunity pursuit (Chikazoe et al., 2014). Although emotion theories differ in considering emotions as discrete entities (Ekman, 1999) versus outputs of continuous dimensions such as valence and arousal (Russell, 2003), they converge in positing that emotions can be evoked by the appraisal of specific survival challenges (Scherer and Moors, 2019). Appraisal theories posit that emotions are elicited and differentiated by a sequence of cognitive evaluations (i.e., appraisals) of the situation (Brosch and Sander, 2013; Lazarus, 1991; Moors et al., 2013; Scherer and Moors, 2019).

The Component Process Model (CPM) conceptualizes emotions as dynamic episodes emerging from multiple components — including motivational changes, autonomic physiological responses, motor expressions, and action tendencies — driven by appraisals of external or internal events (Sander et al., 2005; Scherer, 2009). These appraisal and response components interact to evaluate the evolving meaning of events, and to generate emotional experience as well as adaptive behavior and cognition. In the CPM framework, appraisals correspond to four main objectives, which typically unfold in an automatic and sequential way: i) relevance (i.e., centrality to the self or social reference group), ii) implications (i.e., effects on well-being, immediate or long-term goals), iii) coping potential (i.e., capacity to cope with or to adjust to these consequences of the evaluated object), and iv) normative significance (i.e., significance for personal and for social norms and values) (Sander et al., 2005). Appraisal theories of emotion propose that a series of appraisal processes attribute a value to internal or external events and therefore allow the elicitation of any emotion; such evaluative processes typically include those of novelty, goal relevance, goal congruence, coping potential, compatibility with internal and external standards. The output of such appraisals has been shown to correspond to specific discrete emotions (for a meta-analysis, see Yeo and Ong, 2024).

Neuroimaging studies have provided evidence consistent with the core principles of componential approaches and of appraisal theories including neural dissociability of appraisal processes and emotional responses as well as appraisal-based representations of emotion (Grandjean and Scherer, 2008; Kalisch et al., 2006; Leitao et al., 2020; Mohammadi et al., 2023; Sander et al., 2018; Skerry and Saxe, 2015). For example, novelty detection engages a network centered on the hippocampus (Kumaran and Maguire, 2009) and amygdala (Blackford et al., 2010), concern relevance primarily involves the amygdala (Murray et al., 2023; N’Diaye et al., 2009), goal congruence recruits the anterior cingulate cortex (ACC) (Yee et al., 2021) and ventromedial prefrontal cortex (vmPFC) (Fromer et al., 2019), agency processing is supported by a neural network centered on the temporo-parietal junction (TPJ) (Haggard, 2017), supplementary motor area (SMA), and dorsolateral prefrontal cortex (dlPFC) (Crivelli and Balconi, 2017; Seghezzi et al., 2019) and compatibility with norms and values involves orbitofrontal cortex (OFC) (Zaki et al., 2011), TPJ and dorsomedial prefrontal cortex (dmPFC) (Yomogida et al., 2017). These studies primarily examined appraisal dimensions separately, with limited evidence on their shared neural correlates when multiple dimensions are considered simultaneously.

To the best of our knowledge, only a few neuroimaging studies have included multiple appraisal processes to investigate the brain basis of appraisals and emotions. Skerry and Saxe (2015) found that appraisal processes could account for the neural representations of different emotions in mentalizing regions, including the mPFC, better than basic emotions categories or affective dimensions. Mohammadi et al (2023) found that emotions are brought about through multiple appraisal processes supported by coordinated activity across multiple distributed brain networks. These networks include regions consistently implicated in emotion processing (e.g., amygdala and insula) and those associated with sensorimotor function (e.g., primary somatosensory and motor cortices), social cognition (e.g., precuneus and dmPFC), and attentional control (e.g., dlPFC, ACC, and TPJ) (Mohammadi et al., 2023). However, it remains unclear how these general appraisal networks relate to the neural circuits underlying discrete emotional experiences – particularly epistemic emotions, which have a well acknowledged link to cognitive processes, explaining why these emotions are also called “knowledge emotions” (Chevrier et al., 2019; Erdemli et al., 2025; Pekrun et al., 2017; Silvia, 2026).

While basic emotions support fundamental survival processes such as threat avoidance and aggression, epistemic emotions such as curiosity, interest, and confusion promote exploration and learning (Muis et al., 2015; Pekrun et al., 2017). These emotions arise from individuals’ appraisals of knowledge-related situations, for instance of the discrepancy between new and prior relevant information (Erdemli et al., 2025). Such appraisals involve epistemic incongruity, information novelty, complexity, and self-efficacy/capacity to understand (Muis et al., 2018; Muis et al., 2015). When situations are appraised as incongruent, novel, complex, unexpected, or insufficiently understood, individuals may experience curiosity, confusion, interest, excitement, and awe (Anderson et al., 2020; Murayama et al., 2019; Pekrun et al., 2017; Vogl et al., 2019). In contrast, monotonous situations or those congruent with prior knowledge are more likely associated with boredom (Merrifield and Danckert, 2014). Neuroimaging studies have found that boredom is associated with increased activity in the default mode network (DMN) (Danckert and Merrifield, 2018), and awe with increased activation of the frontoparietal network (FPN) during analytical engagement (van Elk et al., 2019). In turn, epistemic curiosity, which is associated with activation of the reward network (Gruber et al., 2014; Lau et al., 2020), is triggered by prediction errors that elicit appraisals involving the ACC and the lateral prefrontal cortex (LPFC) (Gruber and Ranganath, 2019).

Nonetheless, no study has directly examined the neural mechanisms of multiple appraisal processes underlying epistemic emotions. Instead, appraisal processes have typically been empirically studied in psychology and neuroscience using two methods: 1) experimental manipulation of specific appraisal processes (e.g., novelty or goal-relevance) with measures of various dependent variables such as behavioral responses (e.g., Scherer, 1999), autonomic nervous system responses (Aue and Scherer, 2008), muscular responses (e.g., Gentsch et al., 2015), electroencephalographic markers such as event-related potentials or global field power (e.g., Grandjean and Scherer, 2008; Kalisch et al., 2006; Leitao et al., 2020; Mohammadi et al., 2023; Sander et al., 2018; Skerry and Saxe, 2015), or BOLD responses measured with functional magnetic resonance imaging (fMRI) (e.g., Murray et al., 2023; N’Diaye et al., 2009), or 2) self-report measures of appraisal dimensions across different types of eliciting stimuli (e.g., Israel and Schonbrodt, 2021; Mohammadi et al., 2023; Morgenroth et al., 2026; Scherer and Meuleman, 2013; Smith and Ellsworth, 1985; Brans and Verduyn, 2014; see also see Yeo and Ong, 2024). Traditionally, however, neuroimaging research on emotion have relied on isolated, static stimuli (e.g., emotional figures, facial expression, and words), potentially limiting ecological validity (Shamay-Tsoory and Mendelsohn, 2019) and the examination of dynamic appraisal processes. In contrast, movie-based fMRI provides a naturalistic, dynamic and context-rich approach for studying emotion (Hasson et al., 2004; Mohammadi et al., 2023; Nummenmaa et al., 2023; Saarimaki et al., 2025). By presenting temporally structured emotional events, films reliably evoke dynamic and complex emotional experiences over time. Consequently, movie-fMRI is a promising paradigm for investigating the neural mechanisms underlying appraisals during naturalistic emotional experiences.

### The present study

Here we investigated the brain basis of cognitive appraisals during naturalistic emotional experiences using fMRI, aiming to identify both the common neural circuits shared between appraisal processes and discrete basic and epistemic emotions and the distinctions between them. Participants were scanned with fMRI while watching short movie clips with varying emotional contents. The clips were rated for 12 appraisals and 12 basic and epistemic emotional dimensions by independent participants. We modelled the neural responses measured with fMRI with the ratings of appraisals and emotions to map the appraising brain during naturalistic movie watching. We found that both appraisals and discrete emotions recruited a set of distributed brain regions extending beyond the canonical emotion circuits, encompassing perceptual, motor, and higher-order cognitive systems. These shared activations were more consistent for and better explained by appraisals compared to discrete emotions. Within this network, epistemic emotions elicited less consistent activation than basic emotions, particularly in limbic regions. Our results highlight the functional organization of appraisal and emotional processes under dynamic and naturalistic conditions and suggest that appraisal theories better explain neural responses than models relying only on categorical discrete emotions.

## 2 Methods

### 2.1 Participants

A total of 104 participants were recruited for the fMRI study. The exclusion criteria included a history of neurological or psychiatric disorders, alcohol or substance abuse, BMI under 20 or over 30, current use of psychoactive medications and the standard MRI exclusion criteria. Seven participants were excluded (two due to gradient coil malfunction, two due to structural abnormalities, and three due to visible motion artefacts in the preprocessed fMRI data), resulting in a final sample of 97 participants (50 females; mean age = 31 years; range = 20–57). The study was conducted in accordance with the Helsinki Declaration and was approved by the ethics board of the hospital district of Southwest Finland. An independent sample of 444 healthy volunteers (231 females; mean age = 36 years; range = 18–60) with normal or corrected-to-normal vision were recruited to rate appraisals and subjective discrete emotions elicited by the movie clips. All subjects provided written informed consent and were compensated for their participation.

### 2.2 Stimulus

To investigate the brain responses to emotional stimuli, a well-validated movie dataset was used in the present study (Lahnakoski et al., 2012; Nummenmaa et al., 2023; Santavirta et al., 2023). Specifically, 96 short clips were extracted from Hollywood movies (median duration 11.2 s, range 5.3–28.2 s, total duration 19 min 44 s). During the fMRI data collection, videos were presented in fixed order for all participants, in a single run without breaks. Participants were instructed to watch movie stimuli as if they were viewing a movie at cinema or at home.

### 2.3 Ratings for cognitive appraisals and for discrete emotions

The appraisals used in our study were selected based on the Geneva Appraisal Questionnaire (GAQ, version 3.0) (Scherer, 2001). The selected appraisals included pleasantness, unpleasantness, familiarity, unexpectedness, comprehensibility, complexity, difficulty of coping, acceptability, chance dependency, person agency, intentionality and relevance. Detailed description for appraisal dimensions can be found in supplementary materials (**Table S1**). These appraisals were adapted slightly to ensure their relevance and clarity for the audiovisual stimuli used in our experiment.

Regarding discrete emotions, the six so-called basic emotions were included: sadness, anger, fear, disgust, enjoyment and surprise. Given our focus on appraisal processes that underly the elicitation of epistemic emotions, we included key discrete epistemic emotions considered by current models of emotion, and included both the emotions of “interest” and “curiosity”, in particular to be able to consider their debated relationship (e.g., Tang et al., 2022): confusion, curiosity, boredom, interest, excitement and awe. Altogether, 12 appraisal dimensions, 6 basic emotions and 6 epistemic emotions were evaluated.

The ratings were collected in Gorilla (https://gorilla.sc/), an online behavioral experiment platform. To reduce participant load, the original 33 features were divided into five sets with 6–7 features in each, and each participant was randomly assigned to evaluate one set of features. In addition, the 96 movie clips were randomly assigned into three subsets of 32 movies, and participants were randomly assigned to rate one of the three subsets. For each movie clip, each dimension was ultimately rated by 27–32 participants. See **Figure 1** for the data collection and analysis procedures.

**Figure 1.**
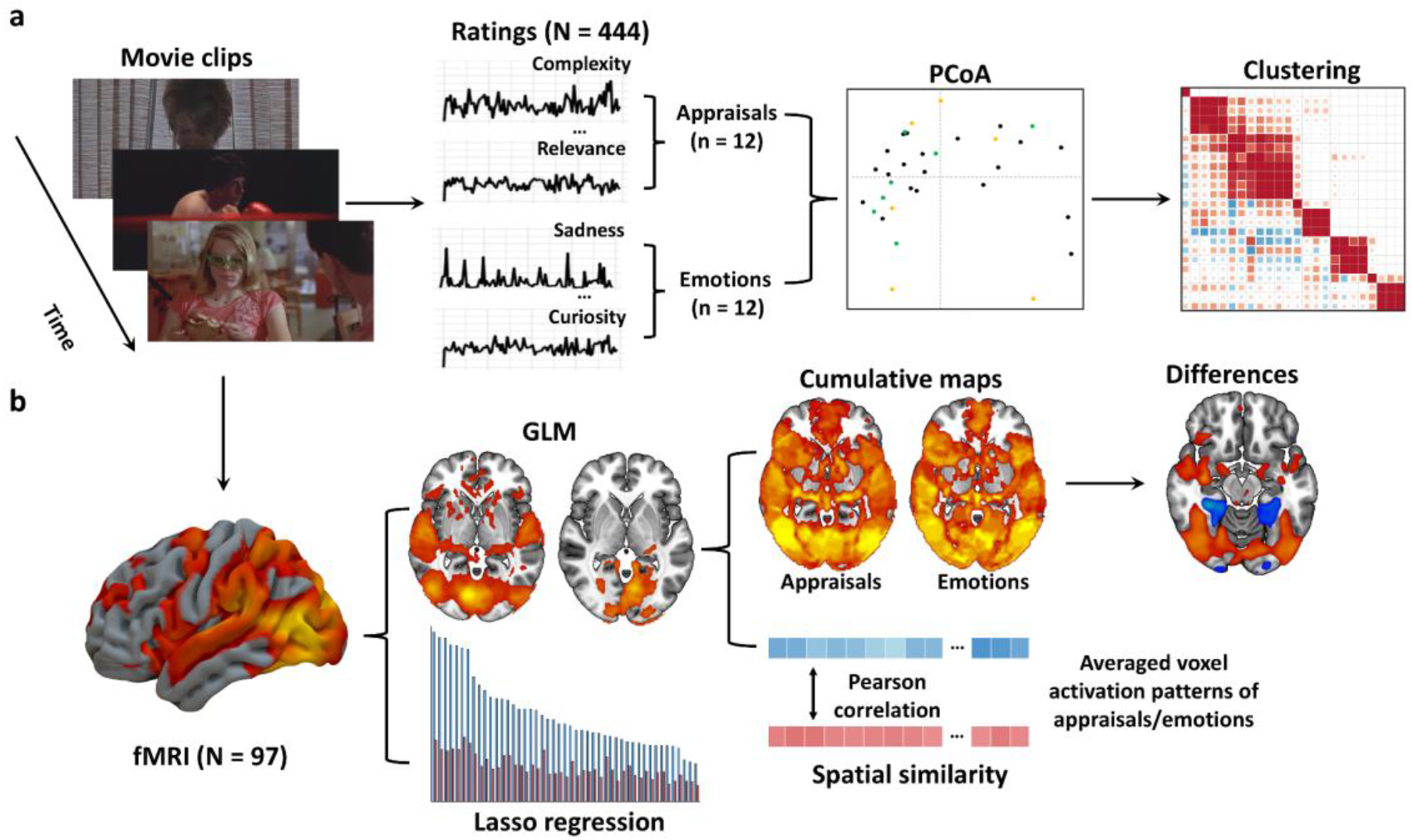
Data acquisition and processing. a) A group of participants (N = 444) were recruited to rate the intensity of 12 appraisals and 12 emotions in the 96 movie clips. The relationship between ratings of appraisals and emotions were then analyzed through principal coordinates analysis (PCoA) and consensus clustering. b) The ratings were subsequently convolved and used as regressors to map the neural representations of appraisals and emotions in an independent fMRI dataset with the same movie stimuli (n = 97). Haemodynamivc responses were compared between appraisals and emotions at both the voxel and regional levels, through cumulative activation, spatial similarity, and Lasso regression analyses.

The ratings were given on an abstract visual analogue scale from 0 (“not at all”) to 100 (“extremely”). See supplementary **Figure S1** for the ratings of appraisals and emotions. To assess the consistency of the ratings between annotators, we calculated the intra-class correlation coefficient (ICC) for each group of participants and video subsets. The ICC2k model was selected since both, the annotators and movie clips, were assumed to represent random samples from their respective populations and videos.

To explore the potential underlying structure of appraisal and emotional features, we applied principal coordinates analysis (PCoA) based on a correlation distance matrix. We then conducted a data-driven consensus clustering analysis using the DiceR package (Chiu and Talhouk, 2018) to examine whether these features exhibited distinct patterns of similarity. The consensus approach ensured a robust and stable clustering solution by performing 1000 independent iterations with random 80% subsamples of the data and over different numbers of clusters (from 2 to 19 clusters).

### 2.4 MRI data acquisition and preprocessing

We collected the MRI data using a Phillips Ingenuity TF PET/MR 3T scanner at Turku PET Centre. Functional volumes were acquired with a T2* - weighted echo - planar imaging (EPI) sequence (TR 2600 ms, TE 30 ms, flip angle 75°, FOV 240 mm, 80 × 80 reconstruction matrix, 62.5 kHz bandwidth, 3.0 mm slice thickness, 45 interleaved axial slices acquired in ascending order without gaps). Structural images were obtained with a T1w sequence (1 mm^3^ resolution, TR 9.8 ms, TE 4.6 ms, flip angle 7°, FOV 256 mm, 256 × 256 reconstruction matrix).

The structural and functional imaging data were preprocessed with fMRIPrep. First and last two functional volumes were discarded to exclude the time points before and after the stimulus. T1w volume was corrected for intensity non-uniformity using N4BiasFieldCorrection (v2.1.0) and skull-stripped using ANTs (v2.1.0) with the OASIS template. Brain surfaces were reconstructed with recon-all FreeSurfer (v6.0.1), and the resulting brain mask was refined by a custom variation of the method to reconcile ANTs-derived and FreeSurfer-derived segmentations of the cortical grey matter of Mindboggle. Spatial normalization to the ICBM 152 Nonlinear Asymmetrical template version 2009c was performed through nonlinear registration with the antsRegistration (ANTs v2.1.0), using brain-extracted versions of both T1w volume and template. Brain tissue segmentation of cerebrospinal fluid, white matter and grey matter was performed on the brain-extracted T1w image using FAST (FSLv5.0.9).

Functional data were preprocessed as follows: slice-timing correction was done using 3dTshift and motion-corrected using MCFLIRT. Next, we co-registered the preprocessed BOLD to the T1w reference image using bbregister for boundary-based registration with six degrees of freedom. All transformations (i.e., motion-correction, coregistration, and spatial normalization) were applied in a single step using antsApplyTransforms (ANTs) and Lanczos interpolation. Spatial smoothing was applied with an isotropic 6-mm Gaussian kernel. ICA-AROMA was performed in MNI space using the standard FSL implementation in a non-aggressive manner to remove motion-related components.

### 2.5 Modelling of the fMRI data

The fMRI data were modelled with SPM12 (Wellcome Trust Center for Imaging, London, UK, https://www.fil.ion.ucl.ac.uk/spm/). To map the brain regions associated with appraisals and emotions, first-level general linear models (GLMs) were constructed separately for each feature of interest. In each GLM, two types of regressors were included: 1) the standardized rating time series of the targeted feature, and 2) the first eight principal components of 14 low-level audiovisual features, including six visual features (luminance, first derivative of luminance, optic flow, differential energy, and spatial energy with two different frequency filters) and eight auditory features (RMS energy, first derivative of RMS energy, zero crossing, spectral centroid, spectral entropy, high frequency energy and roughness) (Santavirta et al., 2023). All regressors were convolved with the canonical hemodynamic response function (HRF) prior to model fitting. Contrast images were generated separately for each participant and subjected to second-level analyses to assess population-level effects. Statistically significant clusters were identified using one sample t-tests (voxel-level threshold *p* < 0.001 with cluster-level family-wise error rate (FWE) correction, *p* < 0.05). At the region of interest (ROI) level, regional effects of each appraisal and emotional dimension were examined using 56 bilateral anatomical ROIs extracted from the AAL2 atlas (Rolls et al., 2015). One sample *t*-tests were conducted on the average β-weights within each ROI to assess the group-level statistical inference, with Bonferroni correction applied for multiple comparisons (*p* < 0.05).

### 2.6 Cumulative maps for cognitive appraisals and emotions

To visualize broad brain networks consistently associated with multiple appraisals and emotions, cumulative maps were generated separately over 12 appraisals and 12 emotions. For each appraisal and emotion, statistically thresholded beta coefficient maps were first binarized (1 = significant effect, 0 = no effect), and then summed across appraisals and emotions. This approach reveals brain networks with the most consistent BOLD responses associated with appraisals or emotions. Similar cumulative analyses were conducted for both basic and epistemic emotions.

### 2.7 Statistical analysis

To test for the overall spatial similarity between appraisals and emotions, we first separately averaged the unthresholded β-coefficient maps for 12 appraisals and 12 emotions, and then calculated the Pearson correlation between the maps.

To explore whether variance in BOLD responses was better explained by appraisal model or discrete emotion model, we conducted Lasso regression at the ROI level. For each participant and ROI, separate Lasso regression models were fitted to predict the BOLD time series from either appraisal-based or emotion-based rating predictors convolved with HRF. A 10-fold cross validation procedure was applied to select the optimal regularization parameter. Predictive performance was quantified using the coefficient of determination (R²), reflecting the proportion of variance in BOLD responses explained by the predictors. For each ROI, differences of predictive performance between appraisals and emotions across participants were assessed using paired sample t-tests. False discovery rate (FDR) correction was used for multiple comparisons.

## 3 Results

### 3.1 Behavioral results

Self-reports for the rated dimensions were mostly reliable, with ICC values mostly over 0.5, and the ICCs were also consistent across the 3 subsets of movies (**Figure 2**). Overall appraisals had the highest ICCs, followed by basic emotions and epistemic emotions. The PCoA revealed a low-dimensional structure underlying the 24 reliable features, with the first two axes together accounting for 71% of the total variance. Based on their distributions in the ordination space, the first axis was interpreted as reflecting affective valence (ranging from unpleasantness to pleasantness), whereas the second axis primarily represented engagement (ranging from boredom to interest) (**Figure 3**). Consensus clustering suggested four stable clusters emerged from the data (**Figure 4**), while the appraisal dimensions of relevance and curiosity remained independent of the four main clusters. The first and largest cluster contained the basic emotions and the basic pleasure-displeasure appraisal dimensions. The second cluster contained interest-related epistemic emotions, while the third cluster consisted of appraisals related to familiarity and comprehensibility and corresponding epistemic emotions. The final cluster focused on intention and agency-related appraisals.

**Figure 2.**
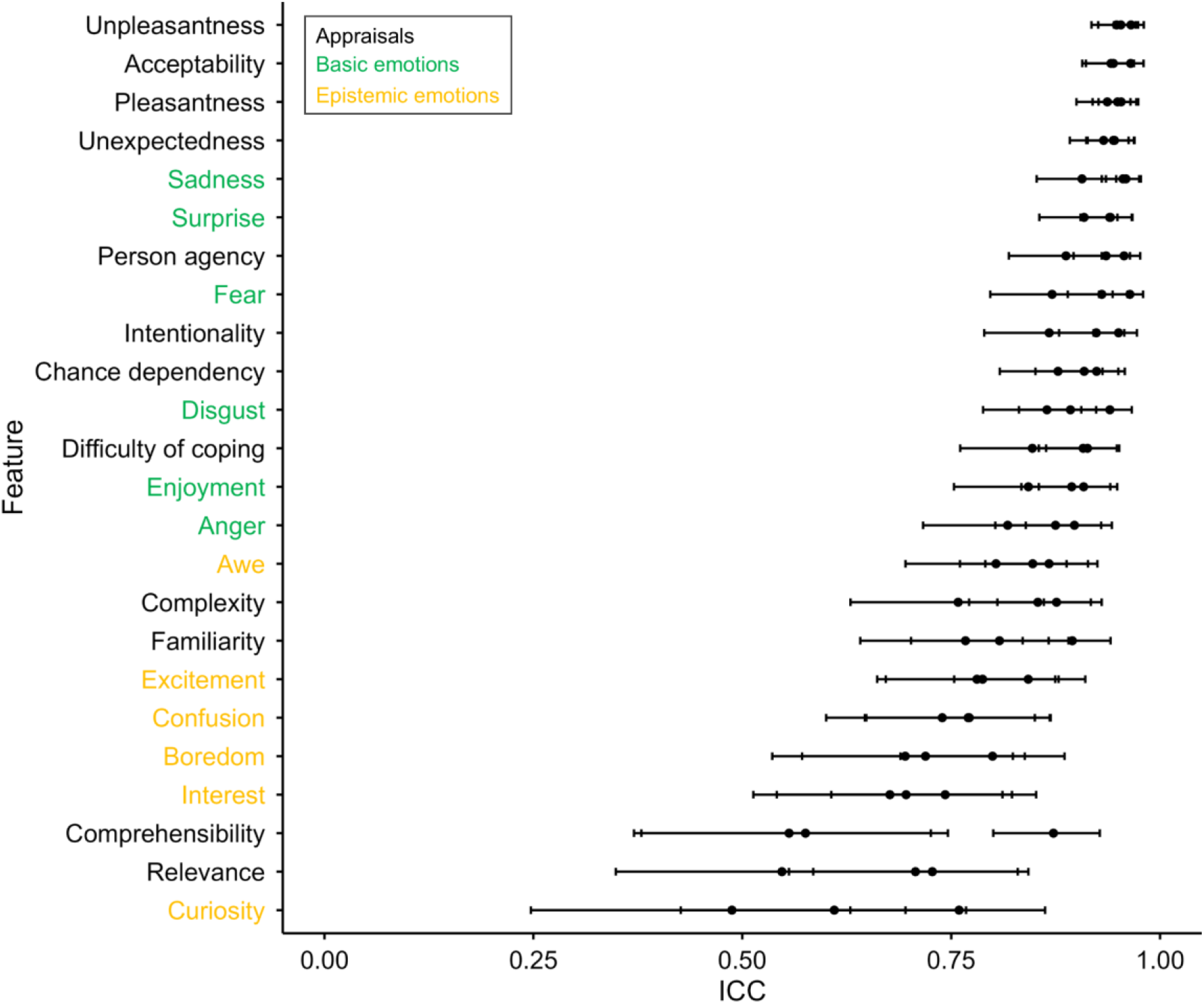
Consistency of appraisal and emotion ratings across annotators. The Y-axis lists dimensions grouped by category (appraisals (n = 12, black), basic emotions (n = 6, green), and epistemic emotions (n = 6, yellow)). The X-axis shows the ICC2k values across participants and stimuli. Individual dots represent reliability estimates obtained from three distinct subsets of movie clips, whereas horizontal black lines indicate the corresponding 95% confidence intervals.

**Figure 3.**
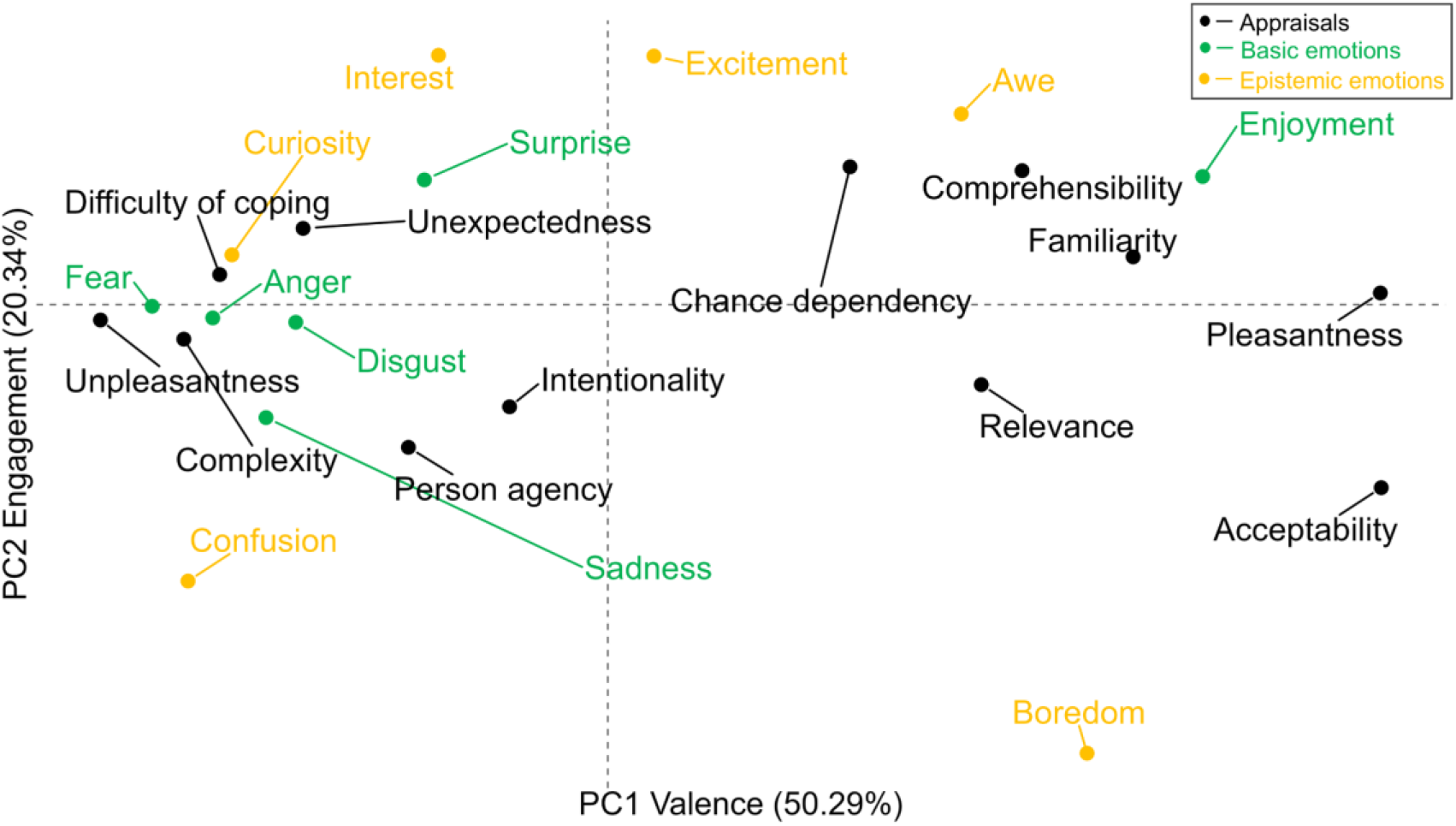
Results of principal coordinate analysis. The plotted first two axes together account for 71% of the total variance (PC1: 50.29%, PC2: 20.34%). Based on their distributions in the ordination space, the first axis was interpreted as reflecting affective valence (ranging from unpleasantness to pleasantness), whereas the second axis primarily represented engagement (ranging from boredom to interest).

**Figure 4.**
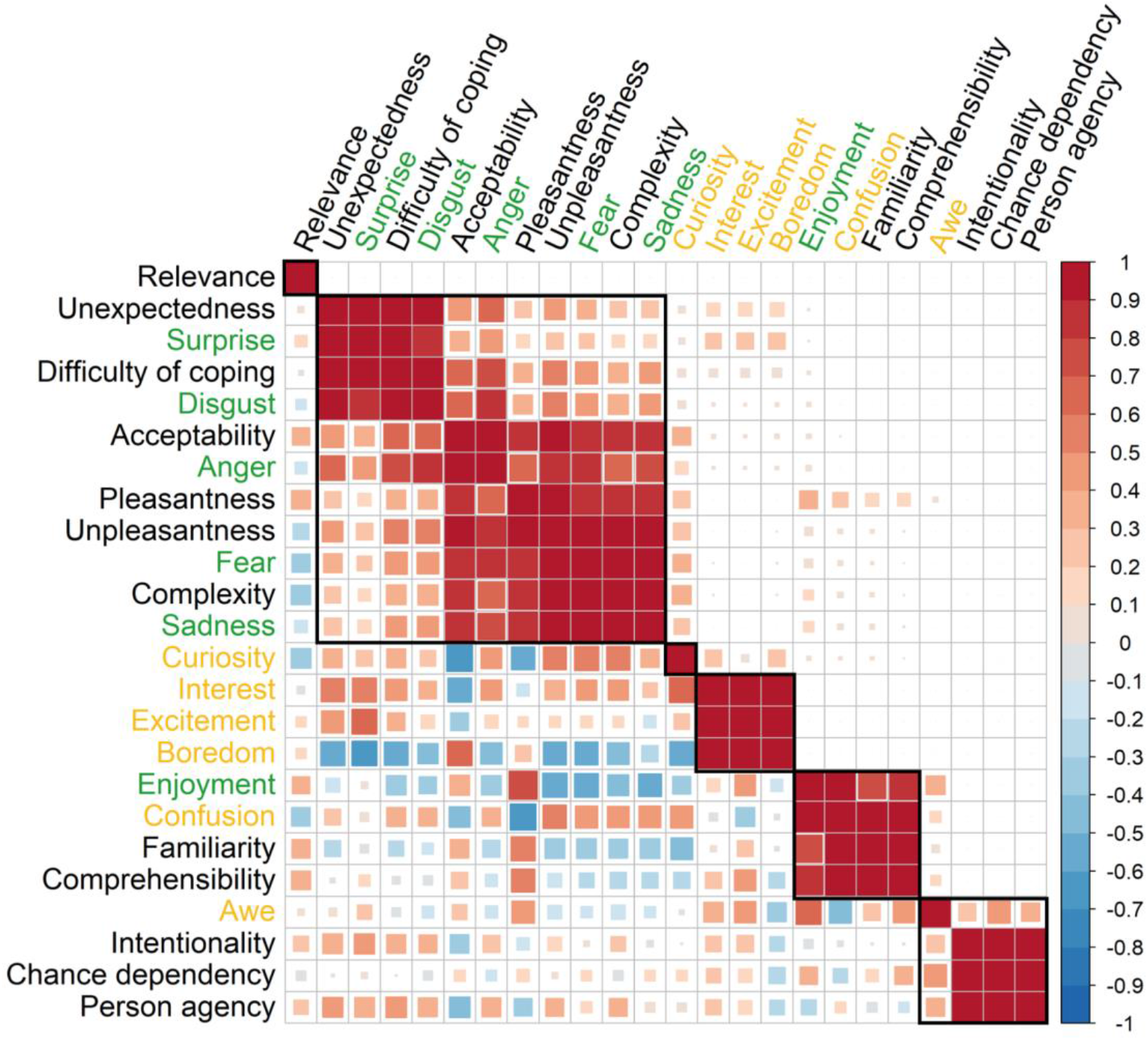
Correlation (lower left) and consensus cluster (upper right) matrix between appraisals (black), basic emotions (green), and epistemic emotions (yellow).

### 3.2 Neural responses to appraisals and discrete emotions

**Figure 5** shows the cumulative brain activation patterns calculated as the sum over statistically thresholded responses for specific dimensions. These maps thus show the brain areas responding consistently to cognitive appraisals and discrete (both basic and epistemic) emotions. Overall, appraisals and emotions activated largely overlapping subcortical and cortical regions. Subcortically, both appraisals and discrete emotions were consistently associated with activation in amygdala, caudate, hippocampus, insula, putamen, and thalamus. Cortically, consistent activation was observed across appraisals and emotion categories in OFC, inferior frontal cortex (IFG), dlPFC, dmPFC, ACC, mid cingulate cortex (MCC), precuneus, SMA, superior temporal gyrus (STG), lateral occipital cortex (LOC) and fusiform gyrus. Despite these substantial overlaps, direct comparison revealed that appraisals were associated with higher cumulative involvement in amygdala, hippocampus, putamen, LOC, STG, dlPFC, dmPFC, SMA, precuneus, IFG, and fusiform gyrus than emotions.

**Figure 5.**
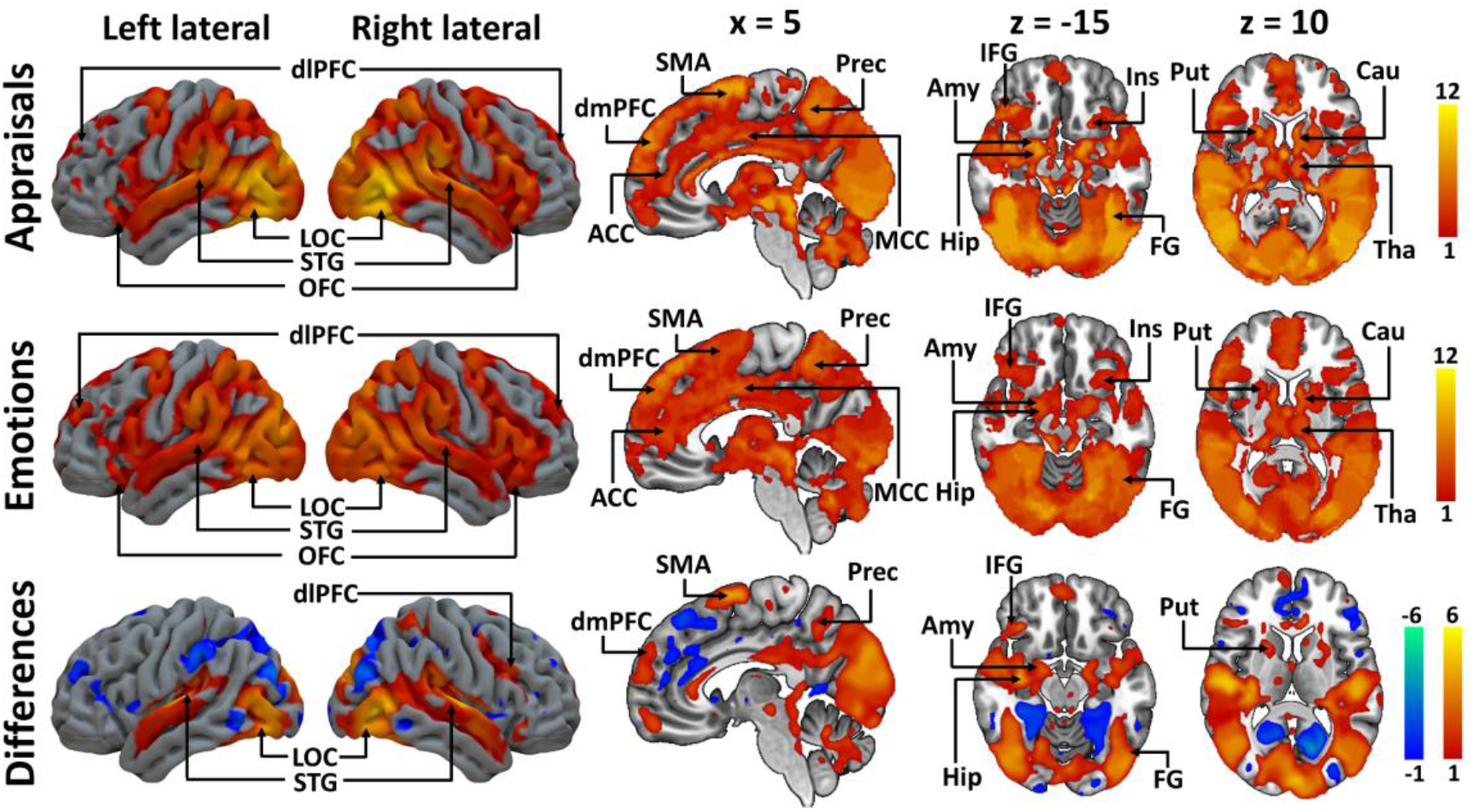
Cumulative activations for appraisals, emotions, and their differences. The color bars indicate the number of appraisals (top row) or discrete emotions (middle row) showing significant associations with the BOLD response (cluster-level FWE corrected at *p* < 0.05, with an initial cluster-forming threshold of *p* < 0.001). The bottom row shows the difference between the two cumulative maps. Hot colors indicate areas where the cumulative map shows more associations with appraisals compared to the emotions while the cool colors indicate the opposite. ACC = anterior cingulate cortex, Amy = amygdala, Cau = caudate, dlPFC = dorsolateral prefrontal cortex, dmPFC = dorsomedial prefrontal cortex, FG = fusiform gyrus, Hip = hippocampus, IFG = inferior frontal cortex, Ins =insula, LOC = lateral occipital cortex, MCC = mid cingulate cortex, OFC = orbitofrontal cortex, Prec = precuneus, Put = putamen, SMA = supplementary motor area, STG = superior temporal gyrus, Tha = thalamus.

The overall spatial correlation between the brain responses for appraisals and discrete (both basic and epistemic) emotions was 0.894 (*p* < 0.001). Moreover, the predictive performance of Lasso models showed that appraisal-based models (mean = 0.076, SD = 0.074) could explain significantly more variance in haemodynamic responses than emotion-based models across most ROIs (mean = 0.052, SD = 0.050) (paired-t test, FDR-corrected *p* < 0.001) (**Figure 6 and Table S2**). A more specific relationship between appraisals and emotions, as well as the regional effects of each appraisal/emotional dimension can be found in supplementary materials (**Figure S2** and **S3**).

**Figure 6.**
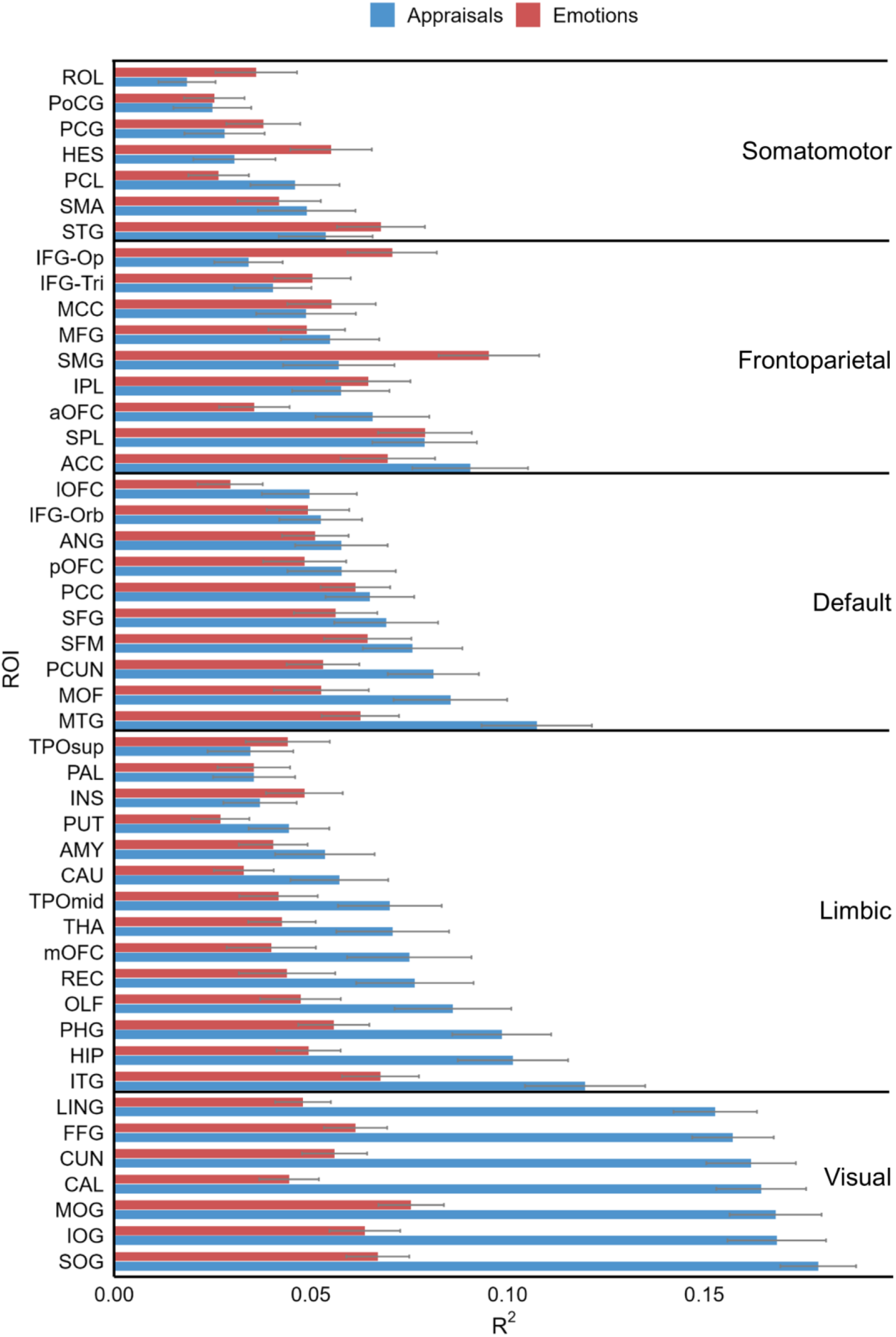
Comparison of predictive performance of Lasso models across 47 ROIs. The bar plot shows the mean R² values for appraisal-based (blue) and emotion-based (red) models, with error bars representing 95% confidence intervals. Across most ROIs, appraisal-based models (mean = 0.076, SD = 0.074) consistently explained more variance in neural activity than emotion-based models (mean = 0.052, SD = 0.050), FDR-corrected *p* < 0.001. See Table S2 for the full name of the regions and statistical results.

### 3.3 Neural responses to basic and epistemic emotions

Finally, to characterize the differences in neural responses between basic and epistemic emotions, we conducted similar cumulative mapping analysis for these emotion types. Cumulative results of basic emotions revealed a widely distributed emotion network (**Figure 7**), including amygdala, caudate, hippocampus, insula, putamen, thalamus, dlPFC, LOC, STG, OFC, dmPFC, ACC, SMA, precuneus, MCC, fusiform gyrus, and IFG. Cumulative maps of epistemic emotions highlighted activation in amygdala, caudate, hippocampus, insula, dlPFC, LOC, STG, dmPFC, ACC, precuneus, MCC, fusiform gyrus, and IFG. Compared to epistemic emotions, basic emotions more consistently engaged amygdala, hippocampus, insula, putamen, thalamus, dlPFC, LOC, STG, OFC, dmPFC, SMA, precuneus, fusiform gyrus, and IFG.

**Figure 7.**
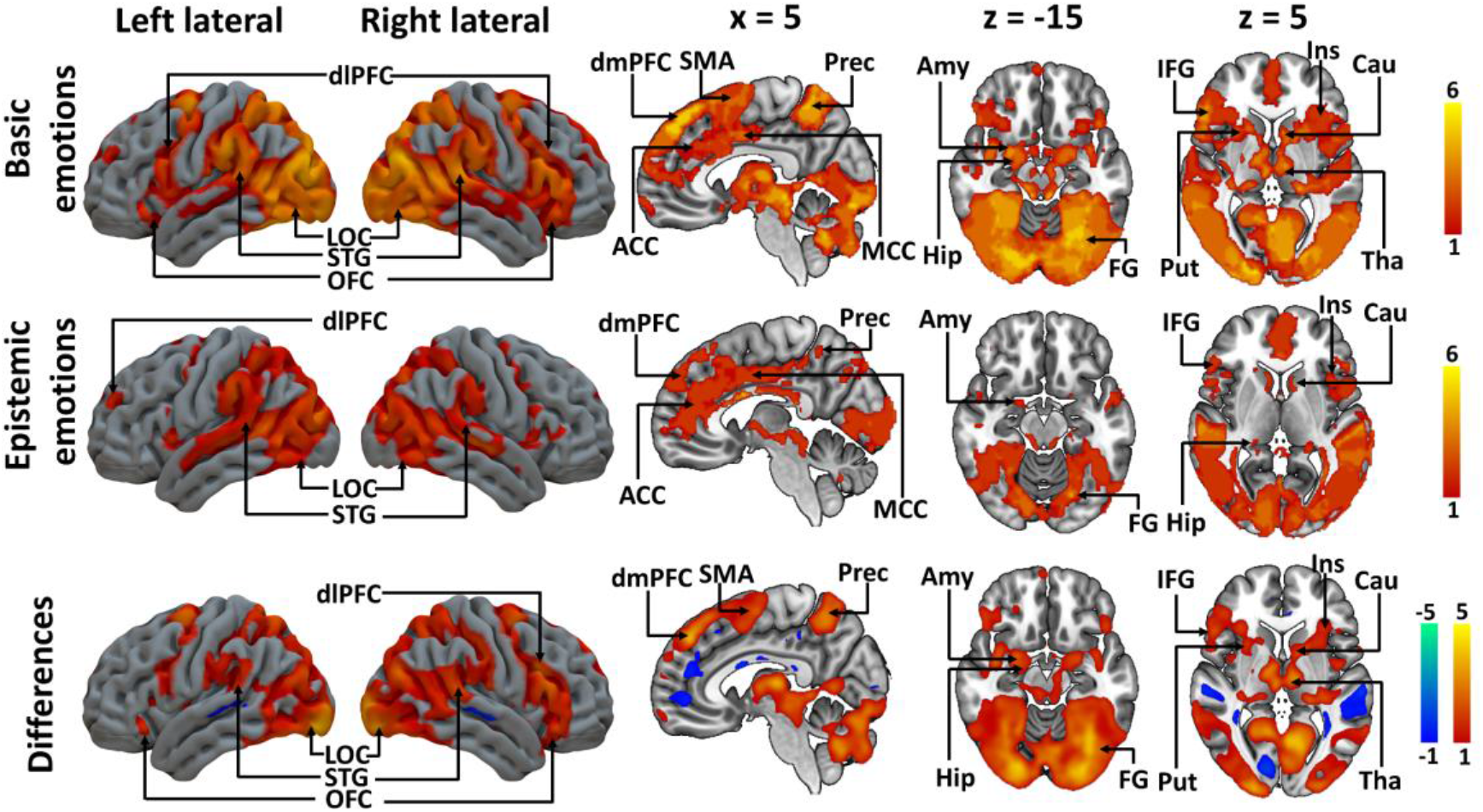
Cumulative activations for basic emotions (top), epistemic emotions (middle row), and their differences (bottom row). The color bars indicate the number of emotions showing significant associations with the BOLD response (cluster-level FWE corrected at *p* < 0.05, with an initial cluster-forming threshold of *p* < 0.001). The bottom row shows the difference between the two cumulative maps. Hot colors indicate areas where the cumulative map shows more associations with basic emotions compared to the epistemic emotions, and cool colors indicate the opposite. ACC = anterior cingulate cortex, Amy = amygdala, Cau = caudate, dlPFC = dorsolateral prefrontal cortex, dmPFC = dorsomedial prefrontal cortex, FG = fusiform gyrus, Hip = hippocampus, IFG = inferior frontal cortex, Ins =insula, LOC = lateral occipital cortex, MCC = mid cingulate cortex, OFC = orbitofrontal cortex, Prec = precuneus, Put = putamen, SMA = supplementary motor area, STG = superior temporal gyrus, Tha = thalamus.

## 4 Discussion

Our main finding was that appraisals as well as discrete emotions recruit a shared large-scale network, including perceptual, motor, limbic, and higher-order cognitive systems. Compared to discrete emotions, appraisals elicited more consistent neural responses across dimensions and accounted for most of the variance in this network. Moreover, neural responses within this network varied across emotion categories: epistemic emotions elicited less consistent activation than basic emotions, particularly in limbic regions. These findings are in line with the proposal that appraisals of events or situations that are of personal significance are an important aspect of emotions (Scherer, 2009).

### Brain basis of appraisals

Appraisals increased activation in multiple perceptual regions (i.e., STG, fusiform gyrus, and LOC). The STG has been shown to be involved in passive detection of novel stimuli (Wong et al., 2026), suggesting a bottom-up orienting response to salient context changes. Notably, the affective salience of novel stimuli can directly enhance the activity in visual regions, such as the fusiform gyrus and LOC, through their dense connectivity with the amygdala (Duncan and Barrett, 2007; Krebs et al., 2009). Such enhanced visual processing may facilitate rapid sensory evaluation, attentional orienting, memory encoding, and assignment of motivational salience (Ranganath and Rainer, 2003). Moreover, these regions are also engaged during social cognitive processing (Santavirta et al., 2023), aligning with evidence that social and affective dimensions interact during appraisal processes in the occipitotemporal cortices (Schacht and Vrticka, 2018; Vrticka et al., 2012). Altogether, the enhanced responses in visual regions are consistent with augmented sensory processing aligning with appraisal models proposing that emotional processing begins with the detection of novel and salient events (Ellsworth, 2013).

Appraisal dimensions were also associated with increased activation in the prefrontal cortex, including the IFG, OFC, dmPFC, dlPFC, ACC and MCC. The IFG has been implicated in attentional orienting and salience detection (Wong et al., 2026). The OFC plays a central role in value representation (Rolls, 2000; Wallis, 2011) and emotion regulation (Hiser and Koenigs, 2018). The dmPFC in turn is associated with social evaluation and inference of others’ intentions (Dixon et al., 2017; Venkatraman et al., 2009). The dLPFC is responsible for the attention allocation (Brosnan and Wiegand, 2017) and belief updating (Huber et al., 2015). The ACC, combined with MCC, contributes to conflict monitoring (Comte et al., 2016; Rushworth and Behrens, 2008) and adjustment of cognitive control (Colin et al., 2025; Gandaux et al., 2025). Together, these regions may form a coordinated network supporting attentional allocation, value representation, and cognitive control during appraisals.

Predictive models of human cognition propose that appraisal-related predictions from prefrontal cortex are unified and integrated with bottom-up sensory inputs in the precuneus, allowing individuals to evaluate current situation using their prior knowledge, and to ultimately shape their emotional experiences (Yazin et al., 2025). Consistent with this account, we observed activation in the precuneus, a central hub of the default mode network that is frequently recruited during self-other evaluation (Cabanis et al., 2013; Legrand and Ruby, 2009) and socioemotional inference (Saarimaki et al., 2025; Skerry and Saxe, 2014). Previous studies demonstrate that the precuneus is involved in the agency appraisal, particularly when individuals attribute the events to external causes (Sperduti et al., 2011). We observed activation in the SMA, which is more involved in self-agency appraisal by generating motor predictions to compare intended actions with actual outcomes (Yomogida et al., 2010; Zapparoli et al., 2020), as well as monitoring actions to reduce discrepancies (Seghezzi et al., 2019). Such activation in the fronto-parietal networks suggests that, while appraisal processes may typically take place automatically, some appraisals may require effortful and higher-level cognitive computations at conceptual level (Scherer, 2009), in particular when participants are requested to explicitly rate the appraisal dimensions like in our study.

We also found extensive appraisal-related activation in the core regions of the affective circuits such as the amygdala, hippocampus, insula, putamen, thalamus, and caudate when appraising the naturalistic movie stimuli. These regions have been associated with a variety of affective processes (Zhang et al., 2025a; Zhang et al., 2025b). Specifically, the amygdala encodes biological and motivational value and plays a key role in emotional processing (Leitao et al., 2022; Mihov et al., 2013). The dorsal striatum, including the caudate and putamen, is involved in reward-related processing (Di Martino et al., 2008; Zheng et al., 2023). The hippocampus supports the encoding and retrieving emotional memories with past experiences (Dimsdale-Zucker et al., 2022; Maurer and Nadel, 2021), the insula is involved in interoceptive awareness (Chong et al., 2017; Craig, 2009), and the thalamus integrates multisensory input to facilitate these processes (Scheliga et al., 2023). The engagement of canonical affective circuits alongside above perceptual and fronto-parietal regions aligns with the notion that appraisals are processed through interacting, hierarchically organized levels: the sensory-motor, schematic, association, and conceptual levels (Scherer, 2009). From this perspective, our data suggest that the emotional responses are not merely outcomes of appraisals, but may also serve as inputs (i.e., monitoring subsystem) to subsequent evaluative processes, including reappraisal, which are well manifested as emotion regulation (Yih et al., 2019; Zhang et al., 2025a). This highlights the dynamic and iterative nature of appraisals, which are continuously updated through interactions with other emotional components, such as subjective feelings.

### Relationships between appraisals and discrete emotion processes

We observed substantial spatial overlap between appraisal and discrete emotion related activity across distributed brain regions, suggesting that evaluative appraisals and discrete emotional processes recruit shared neural substrates. This aligns with prior behavioral evidence demonstrating associations between appraisals and emotions (Conte et al., 2023; Meuleman et al., 2019; Stavraki et al., 2021; Tong and Jia, 2017), as well as findings from neuroimage studies (Skerry and Saxe, 2015). We further found that appraisals accounted for a greater proportion of variance in these brain responses than emotions, suggesting that the activation was primarily driven by appraisals. The dominant role of appraisals is consistent with converging evidence from electroencephalography and fMRI studies (Schacht and Vrticka, 2018; Vrticka et al., 2012), highlighting the importance of appraisal processes in shaping emotional responses. Our study extends previous findings by demonstrating that appraisals engage an extensive network across perceptual, motor, limbic, and higher-order cognitive regions.

Despite this shared large-scale appraisal network, neural responses associated with discrete emotions varied between basic and epistemic emotions. Overall epistemic emotions elicited less consistent activation than basic emotions, particularly in limbic regions. One possible explanation is that different combinations of appraisal dimensions contribute to distinct emotions (Mohammadi et al., 2023; Yeo and Ong, 2024), which may be reflected in broadly distinct patterns between basic *versus* epistemic emotions. Basic emotions, which typically reflect automatic, and evolutionarily conserved affective responses, may engage more prototypical survival-related appraisal profiles (Bach and Dayan, 2017). For example, fear, anger, and sadness are consistently associated with appraisals of unpleasantness, harm, loss, severity, and threat (Yeo and Ong, 2024). In contrast, epistemic emotions rely on a broader range of appraisal processes than basic emotions, as they are more context-specific and are elicited by evaluations of one’s cognitive and knowledge states (Muis et al., 2018). This greater diversity of underlying appraisals could contribute to increased heterogeneity in their neural representations. For epistemic emotions, early appraisals may establish the stimulus relevance with particular engagement of fronto-parietal networks supporting higher-order cognition, such as complex reasoning, memory retrieval, and conceptual integration (Meliss et al., 2024). This pattern is consistent with the CPM’s assumption of a sequential and recursive appraisal process, in which processing may shift toward higher levels when lower-level evaluations are insufficient to resolve the situation (Scherer, 2009).

## 5 Limitations

The passive viewing of movie stimuli likely promoted a predominantly third person perspective and may have limited the extent to which truly subjective emotional experiences potentially attenuating emotional engagement and elicitation of personally relevant experiences. For instance, our framing of the appraisal of “relevance” may not have been appropriate for fiction videos given that we mentioned the “daily life” of the participants. However, previous studies using audiovisual stimuli have shown that even passive viewing or listening can simulate the emotional experiences as reflected by both subjective ratings and synchronization of neural activity (Nummenmaa et al., 2014; Smirnov et al., 2019). Consistent with this, our findings indicate that subjective ratings for appraisals were consistent across participants, supporting the validity of the movie stimuli for eliciting emotional responses. However, future studies using immersive paradigms such as first-person-video-games could further characterize the appraisal processes during situations warranting actions instead of mere passive observation. Importantly, a clear categorization of emotional states as “basic” or “epistemic” is complex and not obvious, and may limit the interpretability of our findings regarding these two emotion “classes”. For instance, previous work highlighted that surprise occupies a boundary position between basic (Ekman et al., 1987) and epistemic emotions (Pekrun et al., 2017). Nevertheless, our results indicate that the overall activation patterns in basic emotions were generally consistent and more widespread than those for the epistemic emotions, indicating that such a separation is conceivable at the neural level, even if further research is needed to reach a firm conclusion. Finally, given that the limited number of discrete emotions is sampling a restricted portion of the affective space, whereas appraisal dimensions capture a wider affective space for more affective states, future studies should recruit more emotions to test the robustness of our findings.

## 6 Conclusions

We identified a distributed large-scale network supporting both appraisals and discrete emotions using emotionally evocative naturalistic stimuli, integrating perceptual, motor, limbic, and higher-order cognitive brain systems. Appraisals play a central role in driving these shared brain activities. Together, these findings suggest that emotion-related brain responses are primarily shaped by appraisal processes, which provides a common computational framework for understanding how different emotions are represented in the brain.

## Supplementary materials

**Figure S1.**
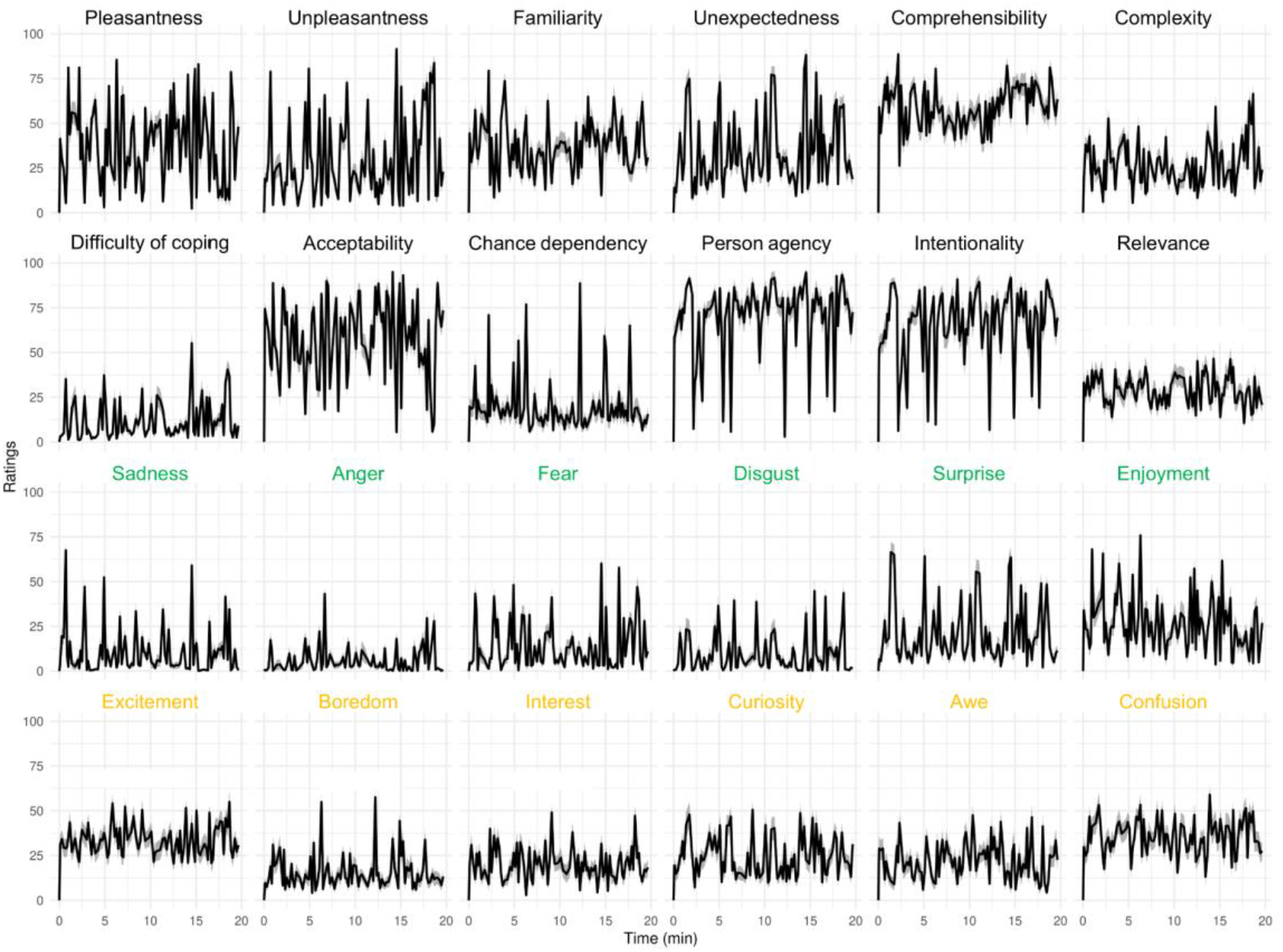
Ratings of 12 appraisals (black), 6 basic emotions (green), and 6 epistemic emotions (yellow).

**Figure S2.**
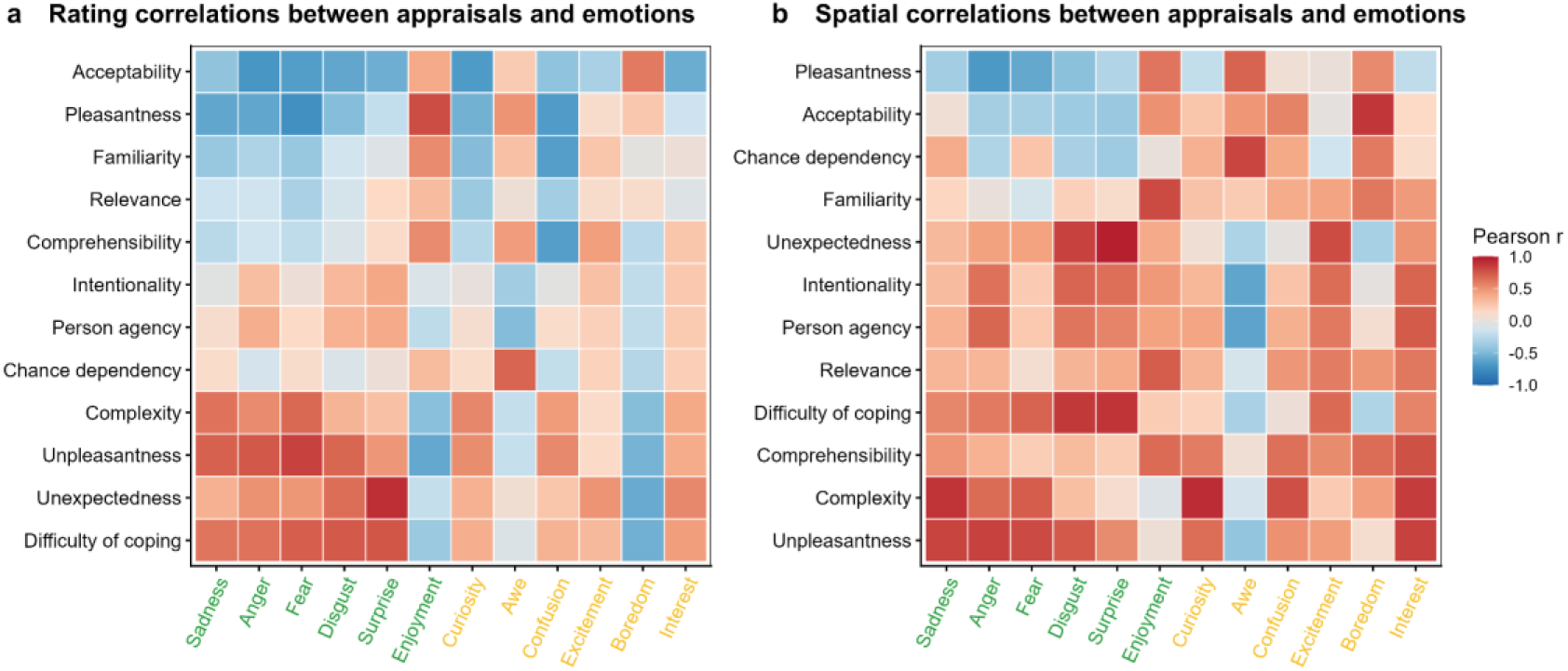
Correlations between appraisals (black) and emotions (basic emotions in green, and epistemic emotions in yellow) computed from ratings (a) and unthresholded β-coefficients (b).

**Figure S3.**
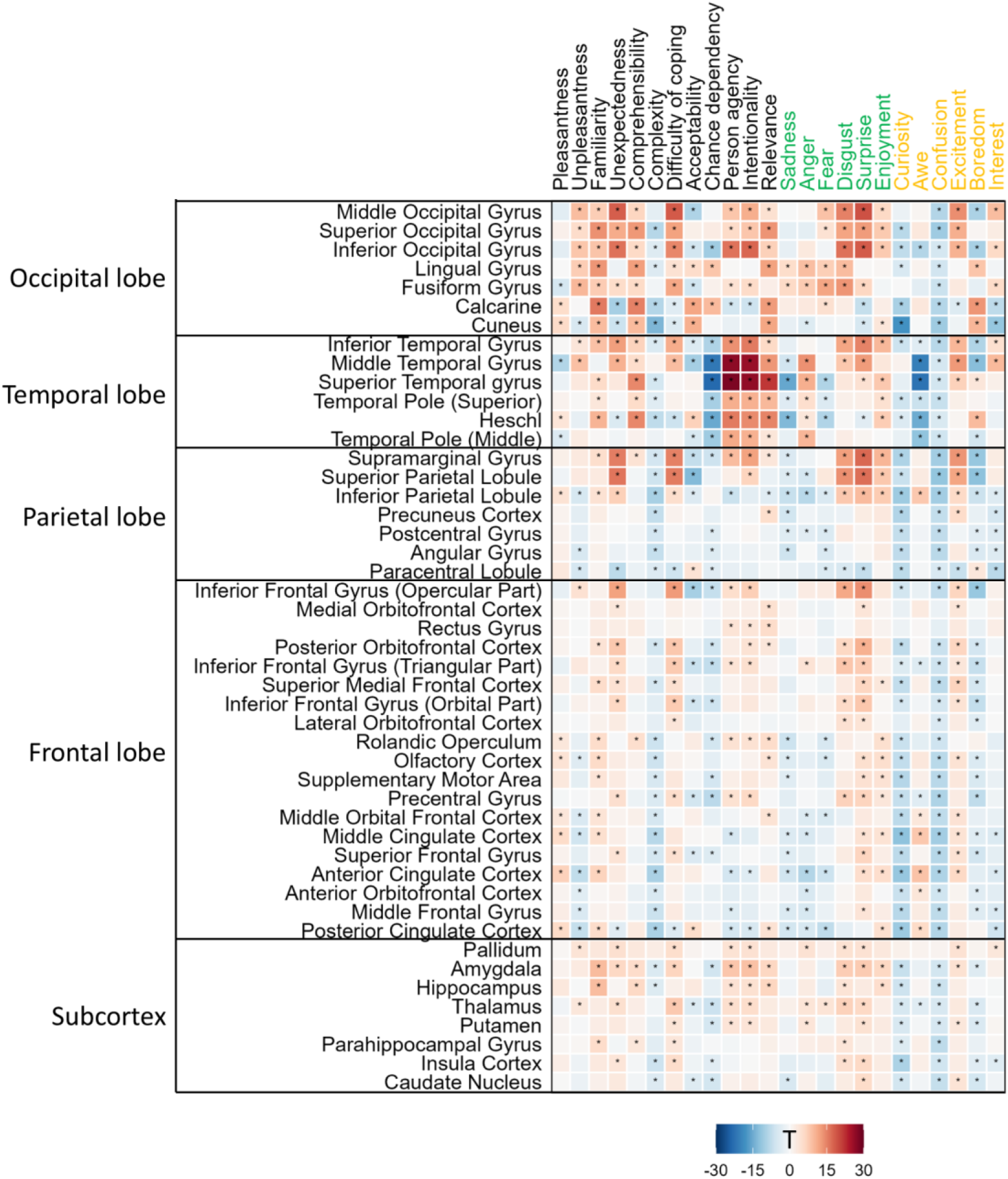
The heatmap indicates the group-level T-values obtained from one-sample t-tests on the average regression coefficients within each ROI for each appraisal/emotional dimension. Statistically significant ROIs (*p* < 0.05, Bonferroni-corrected across ROIs for each dimension independently) are marked with an asterisk.

**Table S1.** Descriptions of appraisal dimensions.

| **Appraisal dimensions** | **Descriptions** |
| --- | --- |
| Pleasantness | "To what extent do you think this situation/event is intrinsically pleasant?" |
| Unpleasantness | "To what extent do you think this situation/event is intrinsically unpleasant?" |
| Familiarity | "To what extent do you think this situation/event is familiar" |
| Unexpectedness | "To what extent do you think this situation/event is unexpected?" |
| Comprehensibility | "To what extent do you think this situation/event is easy to understand?" |
| Complexity | "To what extent do you think this situation/event is complex?" |
| Difficulty of coping | "Too much to handle or hard to cope with" |
| Acceptability | "morally or socially acceptable" |
| Chance dependency | "Consequence of chance, special circumstances, or natural forces" |
| Person agency | "A consequence of behavior of one or more other person(s)" |
| Intentionality | "If so, did these other persons cause the event/situation intentionally" |
| Relevance | "The topics, characters & situations were relevant for my concerns & values in daily life" |

**Table S2.** Brain regions and statistical results presented in Figure 6. Asterisks denote significance levels for a paired t-test between appraisal and emotion models across participants, with FDR correction for multiple comparisons across ROIs. Cohen’s d indicates the corresponding effect size. *FDR corrected *p* < 0.05, \*\**p* < 0.01, \*\*\**p* < 0.001. CI_lower and CI_upper denote the lower and upper bounds of the 95% confidence interval, respectively.

| Full Name | Abbreviation | *t* | *p* | Cohen's *d* | CI_lower | CI_upper |
| --- | --- | --- | --- | --- | --- | --- |
| Rolandic Operculum | ROL | -2.879 | 0.008** | -0.292 | -0.030 | -0.005 |
| Postcentral Gyrus | PoCG | -0.099 | 0.956 | -0.010 | -0.010 | 0.009 |
| Precentral Gyrus | PCG | -1.604 | 0.146 | -0.163 | -0.022 | 0.002 |
| Heschl | HES | -3.15 | 0.003** | -0.320 | -0.040 | -0.009 |
| Paracentral Lobule | PCL | 3.707 | 0.001*** | 0.376 | 0.009 | 0.030 |
| Supplementary Motor Area | SMA | 1.044 | 0.329 | 0.106 | -0.006 | 0.020 |
| Superior Temporal Gyrus | STG | -1.837 | 0.1 | -0.187 | -0.029 | 0.001 |
| Inferior Frontal Gyrus (Opercular Part) | IFG-Op | -6.362 | 0.000*** | -0.646 | -0.048 | -0.025 |
| Inferior Frontal Gyrus (Triangular Part) | IFG-Tri | -1.76 | 0.111 | -0.179 | -0.022 | 0.001 |
| Middle Cingulate Cortex | MCC | -1.049 | 0.329 | -0.107 | -0.019 | 0.006 |
| Middle Frontal Gyrus | MFG | 1.045 | 0.329 | 0.106 | -0.005 | 0.017 |
| Supramarginal Gyrus | SMG | -4.679 | 0.000*** | -0.475 | -0.054 | -0.022 |
| Inferior Parietal Lobule | IPL | -1.101 | 0.319 | -0.112 | -0.020 | 0.006 |
| Anterior Orbitofrontal Cortex | aOFC | 4.034 | 0.000*** | 0.410 | 0.015 | 0.045 |
| Superior Parietal Lobule | SPL | -0.014 | 0.991 | -0.001 | -0.012 | 0.012 |
| Anterior Cingulate Cortex | ACC | 2.739 | 0.011* | 0.278 | 0.006 | 0.036 |
| Lateral Orbitofrontal Cortex | IOFC | 3.335 | 0.002** | 0.339 | 0.008 | 0.032 |
| Inferior Frontal Gyrus (Orbital Part) | IFG-Orb | 0.491 | 0.66 | 0.050 | -0.010 | 0.016 |
| Angular Gyrus | ANG | 1.118 | 0.317 | 0.114 | -0.005 | 0.018 |
| Posterior Orbitofrontal Cortex | pOFC | 1.325 | 0.229 | 0.135 | -0.005 | 0.023 |
| Posterior Cingulate Cortex | PCC | 0.573 | 0.612 | 0.058 | -0.009 | 0.016 |
| Superior Frontal Gyrus | SFG | 1.709 | 0.121 | 0.174 | -0.002 | 0.028 |
| Superior Medial Frontal Cortex | SFM | 1.526 | 0.166 | 0.155 | -0.003 | 0.026 |
| Precuneus | PCUN | 5.261 | 0.000*** | 0.534 | 0.017 | 0.039 |
| Middle Orbital Frontal Cortex | MOF | 4.176 | 0.000*** | 0.424 | 0.017 | 0.048 |
| Middle Temporal Gyrus | MTG | 5.165 | 0.000*** | 0.524 | 0.028 | 0.062 |
| Temporal Pole (Superior) | TPOsup | -1.339 | 0.229 | -0.136 | -0.024 | 0.005 |
| Pallidum | PAL | 0.011 | 0.991 | 0.001 | -0.012 | 0.012 |
| Insular Cortex | INS | -1.815 | 0.102 | -0.184 | -0.024 | 0.001 |
| Putamen | PUT | 3.242 | 0.003** | 0.329 | 0.007 | 0.028 |
| Amygdala | AMY | 2.034 | 0.066 | 0.206 | 0.000 | 0.026 |
| Caudate Nucleus | CAU | 3.945 | 0.000*** | 0.401 | 0.012 | 0.037 |
| Temporal Pole (Middle) | TPOmid | 4.048 | 0.000*** | 0.411 | 0.014 | 0.042 |
| Thalamus | THA | 4.293 | 0.000*** | 0.436 | 0.015 | 0.041 |
| Medial Orbitofrontal Cortex | mOFC | 3.976 | 0.000*** | 0.404 | 0.018 | 0.053 |
| Rectus Gyrus | REC | 3.765 | 0.001*** | 0.382 | 0.015 | 0.050 |
| Olfactory Cortex | OLF | 5.222 | 0.000*** | 0.530 | 0.024 | 0.053 |
| Parahippocampal Gyrus | PHG | 6.861 | 0.000*** | 0.697 | 0.030 | 0.055 |
| Hippocampus | HIP | 8.601 | 0.000*** | 0.873 | 0.040 | 0.064 |
| Inferior Temporal Gyrus | ITG | 6.375 | 0.000*** | 0.647 | 0.036 | 0.068 |
| Lingual Gyrus | LING | 17.991 | 0.000*** | 1.827 | 0.093 | 0.116 |
| Fusiform Gyrus | FFG | 17.255 | 0.000*** | 1.752 | 0.085 | 0.107 |
| Cuneus | CUN | 17.491 | 0.000*** | 1.776 | 0.094 | 0.118 |
| Calcarine | CAL | 19.291 | 0.000*** | 1.959 | 0.108 | 0.132 |
| Middle Occipital Gyrus | MOG | 14.837 | 0.000*** | 1.506 | 0.080 | 0.105 |
| Inferior Occipital Gyrus | IOG | 15.357 | 0.000*** | 1.559 | 0.091 | 0.118 |
| Superior Occipital Gyrus | SOG | 20.17 | 0.000*** | 2.048 | 0.101 | 0.123 |

## Acknowledgements

This work was supported by the Finnish Governmental Research Funding for Turku University Hospital and for the Western Finland collaborative area, Jane and Aatos Erkko Foundation, Gyllenberg’s Stiftelse, European Research Council (ERC Advanced Grant #101141656 to LN), and China Scholarship Council (202408330143).

